# *Mycoplasma gallisepticum* uses itaconate-associated mitochondrial inhibition to suppress host immunometabolism

**DOI:** 10.64898/2026.08.12.744485

**Authors:** Soren Z Coulson, Reihane Eric, Chidambaram Ramanathan, Katie M Talbott, Francis E Tillman, Anna Perez-Umphrey, Truc Chi Thi Pham, Paul S Simone, Brandt D Pence, James S Adelman, Yufeng Zhang

## Abstract

Many pathogens actively suppress early host immune responses to enhance their fitness. Mitochondria function as key regulators of immune activation, yet whether pathogens suppress host immunity by manipulating mitochondrial metabolism *in vivo* remains largely unknown. During an innate immune response, the metabolite itaconate increases in abundance and acts as an immunomodulator, due to its inhibition of succinate dehydrogenase (SDH), a key mitochondrial regulator of cellular immunity. We hypothesized that *Mycoplasma gallisepticum* (MG), a recently emerged pathogen of wild songbirds, most notably house finches (*Haemorhous mexicanus*), suppresses early host immune responses by limiting SDH-dependent immune activation via itaconate. We tested these hypotheses using experimental 3-day infection of finches with heat-killed MG, live MG or pharmacological elevation of the SDH inhibitor itaconate. Following inoculation, we quantified intracellular itaconate and mitochondrial respiratory function in peripheral blood mononuclear cells (PBMCs) and pro-inflammatory cytokine gene expression in erythrocytes, in addition to infected tissues (trachea and conjunctiva). Heat-killed MG increased SDH-dependent mitochondrial respiration in PBMCs and cytokine gene expression in erythrocytes, but live MG did not show these increases, but revealed increased itaconate accumulation in PBMCs. Dimethyl itaconate administration reproduced the suppressed metabolic and immune phenotype in blood cells observed with live MG, suggesting an itaconate-associated mechanism. In contrast, live MG increased mitochondrial respiration and gene expression levels of cytokines in eyelid conjunctiva, whereas other treatments did not. These findings indicate that MG suppresses host metabolic and cytokine signaling in systemically circulating immune cells through a mechanism consistent with itaconate-mediated inhibition of SDH-dependent mitochondrial respiration, while still inducing an inflammatory response at the site of infection. Our data suggest that MG, like other pathogens, can commandeer host immunometabolic pathways during infection to their benefit and that mitochondria are a key site of competition between host and pathogen.

## Introduction

Successful pathogens frequently employ strategies that blunt the effectiveness of host defenses. These strategies are highly variable and can include active interference in diverse host immune processes [1–3]. Immune cell metabolic reprogramming has emerged as an essential component of innate immune activation [4–6] and these metabolic responses may be targets of subversion by pathogens [7–11]. Increasing evidence demonstrates that innate immune activation depends on mitochondrial metabolic reprogramming [4,12] and that mitochondria are often targeted by pathogenic bacteria [11,13] yet the metabolic mechanisms underlying pathogen manipulation of host mitochondria require additional investigation. *Mycoplasma gallisepticum* (MG) is an emerging bacterial pathogen in birds that causes respiratory disease in domestic poultry [14] and conjunctivitis in the wild songbird, house finches (*Haemorhous mexicanus*) [15]. MG emerged as a major pathogen in house finches following a host species transfer from domestic poultry in the 1990s [16] and has since spread throughout the house finch range across most of North America [17,18]. MG has been hypothesized to have immuno- and metabolic suppressive properties, such as reducing host lymphocyte activation and altered cytokine responses [19–22], although the underlying mechanisms remain unresolved. Immunometabolic pathways are highly conserved among host taxa [23], so emerging pathogens may target these pathways. Therefore, identifying the pathways targeted by MG may provide broader insight into how bacterial pathogens manipulate conserved immunometabolic processes.

Mitochondria play a significant role in mediating the immune responses to pathogens, acting as a nexus of signaling and ATP production which are requisite for immune cell activation and cytokine production [12,24,25]. Therefore, suppression of mitochondrial metabolism presents a plausible mechanism by which pathogens may impair host immune function [26–29]. MG disrupts host mitochondrial physiology, including mitochondrial swelling in immune tissues [30,31] and decreased succinate dehydrogenase (SDH)-mediated respiration in liver mitochondria [32,33]. SDH has gained recognition as a critical regulator of the immune response due to its role in linking mitochondrial respiration and TCA cycle with inflammatory signaling [34,35], so the apparent inhibition of SDH during MG infection may reflect a general suppression of host immune response in house finches. However, whether this MG-induced SDH inhibition occurs in immune cells and tissues at the site of infection, in addition to underlying mechanisms for this inhibition, remain unclear.

In mitochondria, tricarboxylic acid (TCA) cycle metabolites not only serve as intermediates in macronutrient metabolism, but also function as important signaling molecules involved in a wide range of cellular processes, particularly immune regulation [4]. Among these immunometabolites, itaconate, succinate, and fumarate (the so-called “gang of three”) are all closely linked to SDH and play important roles in regulating immune effector functions [34]. As both an inflammatory signaling molecule and an inhibitor of SDH, itaconate functions predominantly as an endogenous immunosuppressive regulator of host immunity during infection [36–38]. Infections caused by other *Mycoplasma* species, including *M. arginini* and *M. pneumoniae*, have been shown to induce increased itaconate synthesis in immune cells, which has been associated with reduced neutrophil bactericidal activity and increased infection severity [39,40]. Because itaconate is both a potent endogenous inhibitor of SDH and is induced by infections caused by other Mycoplasma species, it represents a particularly strong candidate mechanism linking MG infection to SDH suppression. Together, these observations raise the possibility that MG infection may induce alterations in host itaconate signaling that contribute to suppression of immune metabolic activation during the early stages of infection.

Here, we tested whether MG suppresses host immunometabolism by inhibiting mitochondrial respiration in immune cells and infected tissues, and whether this phenotype resembles pharmacological elevation of itaconate. We tested the hypotheses that 1) MG infection would globally suppress mitochondrial respiration and immune markers in the host and that 2) MG infection would suppress mitochondrial respiration via itaconate. We predict that 1) MG infection will decrease mitochondrial respiration and pro-inflammatory cytokine expression in tissues at the site of infection and in circulating immune cells and that 2) this phenotype will mimic mechanisms of itaconate-mediated immunometabolic suppression. Our findings have the potential to identify immunometabolic suppression as a previously unrecognized mechanism of MG pathogenesis. This study provides new insight into how bacterial pathogens manipulate host mitochondrial metabolism to evade immune defenses, highlighting immunometabolism as a critical but underappreciated interface in host–pathogen interactions.

## Methods

### Experimental animals

All experimental procedures were approved by the University of Memphis Institutional Animal Care and Use Committee (CY25-013). All birds were handled and captured under permits from U.S. Fish and Wildlife Service (MB154804-0), Virginia Department of Game and Inland Fisheries (066646) and Tennessee Wildlife Resources Agency (Import Permit 39263697, Scientific Collecting Permit 2252). Wild house finches were captured (Montgomery County, VA, USA) using basket trap and mist nets, then brought into captivity at the University of Memphis (Memphis, TN, USA) animal care facility. Birds included in these experiments had been part of a prior infection experiment testing how photoperiod and prior exposure to MG impacted both infection severity and reproductive investment (Talbott et al, *in review*). However, most (14 out of 24) animals used in the experiment reported here had not been exposed to MG during prior work (i.e. had served as controls). Birds that had been previously treated with MG (10 out of 24) had recovered from infection for c. 14-15 weeks, with one individual recovered for c. 22 weeks and were evenly distributed across current treatment groups. Prior to the current experiments, birds were kept on 12L:12D photoperiod in flight cages (76 cm x 46 cm x 46 cm) for and given *ad libitum* food and water. To ensure that birds were not actively infected at the start of experiments, we measured plasma MG antibodies using a commercially available kit (IDEXX Laboratories Inc., Westbrook, ME). All birds were free of symptoms prior to beginning experiments, and MG antibody levels were lower than 0.061 OD (optical density) seropositivity threshold cutoff [41,42]. We randomly divided birds into four experimental groups (n = 6). We used the VA94 strain of MG and diluted in Frey’s medium [43] to 7.5 x 10^6^ color-changing units (CCU) mL^-1^. Previous work from our lab groups has shown that hepatic mitochondrial respiration decreases following inoculation of this strain and dosage of MG [33]. We bilaterally inoculated all birds in the palpebral conjunctiva of the eye with 20 µL of MG via pipette, while control birds were inoculated with Frey’s medium, as performed previously [33]. We inoculated the third group of birds with heat-killed MG at the same dosage and volume, which we prepared by incubating MG at 65°C for 45 minutes. We repeated the heat-killed MG inoculation for the fourth group of birds but also injected dimethyl itaconate (a cell-permeable esterified itaconate derivative that has similar anti-inflammatory effects as itaconate [44]) at 800 mg kg^-1^ intraperitoneally (IP). Although dimethyl-itaconate may not fully recapitulate endogenous itaconate signaling, it is a widely used pharmacological tool for investigating itaconate-mediated metabolic regulation [45,46]. Our pilot experiments indicated that PBMCs intracellular itaconate concentration increased roughly three-fold following dimethyl itaconate injection, but was similar between dosages of 400, 800 and 1600 mg kg^-1^ (Fig. S1).

We administered each inoculation and injection at 0, 24 and 48 hours and sampled the birds 72 hours after beginning the experiment. We selected this time point because house finches typically begin to exhibit clinical signs of infection at this stage [47] and our prior studies demonstrated alterations in isolated hepatic mitochondrial function three days post-inoculation [33]. We euthanized the birds via decapitation and collected blood from the neck into a conical tube, which we used to isolate PBMCs for mitochondrial assays or itaconate quantification. We also dissected the trachea and upper and lower eyelid conjunctiva from one eye for immediate use in mitochondrial assays. The eyelid conjunctiva from the other eye was snap-frozen in liquid nitrogen and stored at −80°C until qPCR analysis.

### PBMC isolation

We isolated PBMCs from whole blood using Cytiva Ficoll-Paque™ PLUS media (Cat# 17144002; Cytiva, Marlborough, MA, USA) density-gradient centrifugation according to manufacturer protocol. After Isolation, PBMCs were used for respiration measurements. Due to limitations in PBMC yield, we also isolated PBMCs from a separate cohort of birds (n = 3-4) which were then counted via hemocytometer, snap-frozen in liquid nitrogen and stored at −80°C before measurement of intracellular itaconate via mass spectrometry (see following section).

### Itaconate quantification

We measured itaconate concentration in PBMCs via mass spectrometry analyses as previously described [48]. First, we lysed the PBMCs with the addition of 1 mL of ice-cold (8:2) methanol:H_2_O to each cell pellet, followed by vortexing, sonication, then centrifugation at 15,000 g for 15 minutes at 6°C. We aspirated 450 µL of the resulting supernatant and dried it down in a glass vial, using N_2_ gas. Next, we added 100 µL of 250 mM 3-NPH (dissolved in 1:1 acetonitrile:H_2_O) and 200 µL of 143.5 mM EDC-HCl (dissolved in 1:1 acetonitrile:H_2_O with 6% pyridine (v/v)), vortexed and incubated at room temperature for two hours. Following this step, we added 500 µL chloroform and 100 µL of ice-cold 3N HCl, mixed by vortexing and waited for complete layer separation. We then aspirated 450 µL of the lower organic layer into a 1.2 mL Max Recovery glass vial (Microsolv, Greater Wilmington, NC, USA), which we dried with N_2_ gas. We reconstituted the resulting powder in (0.1:80:20) NH_4_OH:H_2_O:acetonitrile, sonicated for 1 minute and passed through a 0.45 µm Thompson filter vial before UPLC MS/MS analyses.

We used an Acquity UPLC system with Quattro micro triple quadrupole mass spectrometer, outfitted with Waters’ UPLC BEH C18 column (1.7 µm, 2.1 x 150 mm) maintained at 50°C. The mobile phase was a mixture of 0.1% formic acid in H_2_O (A) and 0.1% formic acid in acetonitrile (B). Our gradient elution used 98% A for 0.75 minutes, ramp to 80% until 3 minutes, ramp to 50% until 3.75 minutes, isocratic until 4.2 minutes then ramp to 98% at 4.35 minutes at a constant flow rate of 0.45 mL min^-1^. We used a standard curve to calculate PBMC itaconate abundance. We dissolved itaconic acid in (0.1:80:20) NH_4_OH:H_2_O:acetonitrile, sonicated for 1 minute and diluted to 200, 100, 50, 25 and 12.5 ppb standards. Injection volume was 10-15 µL.

### PBMC respiration measurements

To investigate PBMC respiration, we counted and seeded the PBMCs on the Agilent Seahorse XF cell culture plates at 250K cells/well with Seahorse XF RPMI medium. In our first run, we measured SDH-driven oxygen consumption rate (OCR) in PBMCs following permeabilization with digitonin (15 µg mL^-1^). Pilot experiments showed minimal loss of cytochrome c from PBMCs following digitonin digestion (Fig. S2), indicating that mitochondrial integrity was not compromised. Following permeabilization, we stimulated complex II with succinate (10 mM) and rotenone (0.5 µM) (state 2), followed by stimulation of oxidative phosphorylation with ADP (5 mM) (state 3). Next, we inhibited ATP synthase with oligomycin (1 µg mL^-1^) to measure leak OCR (state 4o). State 2 and state 4o OCR correspond to mitochondrial proton leak, while state 3 OCR corresponds to capacity for ATP synthesis via oxidative phosphorylation. After, we added antimycin A (0.5 µM) to inhibit the electron transport system and measure non-mitochondrial OCR, which we subtracted from all other OCR measurements.

In a separate run, we performed the Cell Mito Stress Test (Cat# 103015-100; Agilent, Santa Clara, CA, USA) in PBMCs following manufacturer’s instruction [49], wherein we measured OCR and ECAR first in the absence of any added substrates or inhibitors (Baseline), then following oligomycin addition (2 µg mL^-1^) to induce leak OCR and glycolytic capacity. Next, we added FCCP (2 µM) to induce maximal OCR, followed by rotenone (0.5 µM) and antimycin A (0.5 µM) to inhibit the mitochondrial electron transport system so that we could measure non-mitochondrial OCR. We subtracted non-mitochondrial OCR from all preceding OCR measurements.

### Tissues respiration measurements

After dissecting and cleaning the trachea and eyelid conjunctiva of feathers and contaminating tissues we permeabilized these tissues, using methods described previously [50]. We digested each tissue in isolation solution (in mM: 2.77 CaK_2_EGTA, 7.23 K_2_EGTA, 20 imidazole, 20 taurine, 49 K-MES, 3 K_2_HPO_4_, 9.5 MgCl_2_, 5.7 ATP, 15 phosphocreatine, 0.001 leupeptin, pH 7.1) with 50 µg mL^-1^ saponin on ice and with gentle mixing for 20 minutes. We then washed each tissue by transferring to ice-cold respiratory medium (in mM: 0.5 EGTA, 3 MgCl_2_ 6H_2_O, 20 taurine, 10 KH_2_PO_4_, 20 HEPES, 60 K-lactobionate, 110 mannitol, 0.3 dithiothreitol; 1 mg mL^-1^ BSA, pH 7.1) with gentle mixing for 5 minutes. We repeated the washing step two additional times. Pilot experiments showed minimal loss of cytochrome c from either tissue following saponin digestion (Fig. S2), indicating that mitochondrial integrity was not compromised. We then dabbed each tissue sample dry using a tissue and measured wet mass, which we used to standardize our OCR measurements. Trachea mass used was 4.7±0.1 mg and eyelid conjunctiva mass was 3.3±0.2 mg. We then placed the tissues into an Oxygraph-2k (Oroboros Instruments, Innsbruck, Austria) for OCR measurements.

We calibrated the Oxygraph-2k daily with air equilibration and at anoxia. We measured OCR in 0.5 mL of respiratory medium set to 40°C. We first measured complex I OCR in the presence of glutamate (10mM) and malate (5mM) (State 2) and then with ADP (10mM) (State 3). Next, we inhibited complex I with addition of rotenone (1.6 µM), and stimulated OCR through complex II with the addition of succinate (20mM). After, we added oligomycin (0.08 µg mL^-1^) to inhibit ATP synthase and reveal OCR caused by mitochondrial H^+^ leak (State 4o). At the end of the experiment, we added antimycin A (5 µM) to completely inhibit mitochondrial OCR, allowing us to measure non-mitochondrial OCR. We corrected all mitochondrial OCR values by subtracting non-mitochondrial OCR and instrumental OCR.

### Quantitative PCR

We assessed the expression of immune markers in the eyelid conjunctiva, targeting 7 genes that have previously been shown to be significantly upregulated following MG inoculation in house finches: *IL1B*, *IL6*, *IL8L*, *IL10*, *IL18*, *TGFB2* and *TNFSF15* [51]. We also measured the expression of 3 reference genes: *28SrRNA*, *GAPDH* and *ACTB*. We also measured the expression of the same genes in blood cells. Due to limitations in PBMC yield, we instead used erythrocytes, which in birds are nucleated and contribute to immune function [52,53]. We isolated RNA by first homogenizing one eyelid conjunctiva or 50 µL of erythrocytes in 200 or 250 µL of TRIzol, respectively, using 0.2 g zirconium oxide beads in a BeadBlaster 24R (Benchmark Scientific) at 4°C (4000 rpm, two 30 s cycles). We then used these homogenates directly with a commercial kit (Zymo Research D7003) to isolate and purify RNA, following manufacturer instructions, which included on-column DNase treatment. We quantified RNA purity and concentration using a Nanodrop Lite (Thermo Scientific).

We reverse-transcribed 0.2 μg of RNA into cDNA using a reverse transcriptase kit (Thermo Fisher 4374966) and thermal cycler (SimpliAmp; Thermo Fisher) set to 25°C for 10 minutes, 37°C for 2 hours, then 85°C for 5 minutes. We diluted the cDNA 5-fold with nuclease-free water prior to qPCR. We used the same primers as those used previously studying the transcriptional response to MG infection in house finches [51], prepared by Integrated DNA Technologies (Morrisville, NC, USA) (Table S1). qPCR reactions were 15 μL in total and were comprised of SYBR Green master mix (Abclonal RK21203), 0.2 μM gene-specific primers, and included 5 µL of cDNA. We plated reactions in a 384-well plate in duplicate for each gene. We also plated 10 µL master mix plus 5 µL of water as a negative control. We monitored DNA amplification using a QuantStudio 6 Flex (Thermo Fisher). At the hold stage, plates were incubated at 50°C for 2 minutes, followed by 95°C for 10 minutes. In the PCR stage, plates were cycled 40 times between 95°C for 15 seconds and 60°C for 1 minute. In the melt curve stage, plates were cycled between 95°C for 15 seconds, 60°C for 1 minute and back to 95°C for 15 seconds. We analyzed the qPCR data with QuantStudio software (version 1.3), which automatically determined threshold ΔRn and baseline values. We averaged Ct values across replicate wells and used 28S rRNA as a reference gene, as used previously [51]. We calculated ΔCt values for each sample by subtracting the Ct value of the reference gene (28S rRNA) from the Ct value of each target gene. We then calculated ΔΔCt values by subtracting the mean ΔCt of the media control group from the ΔCt of each corresponding sample. We express differences in gene expression as fold change, which we calculated as 2^-ΔΔCt^, such that values greater than 1 indicate upregulation and values less than 1 indicate downregulation relative to the media control group.

### Statistical analyses

We analyzed and plotted all data within Prism version 11.0.0 (GraphPad Software, Boston, MA, USA). We assumed normality in our datasets but tested for equal variance using the Brown-Forsythe test, with a significance cut-off of p < 0.05. For comparisons where variance was equal among groups, we compared means using 1-way ANOVA, with Tukey HSD as a post-hoc test. For comparisons where variance was not equal among groups, we compared means using Welch’s ANOVA, with Dunnett’s T3 multiple comparisons test for post hoc analyses. We determined significant differences to be p < 0.05.

## Results

### Itaconate increases in immune cells during MG infection

To investigate the role of itaconate in MG infection, we first measured itaconate levels in PBMCs following MG inoculation (Fig. 1). Intracellular itaconate levels varied among inoculations (F_2,7_ = 9.2, p = 0.01); live MG infection increased intracellular itaconate accumulation by approximately 4-fold relative to media controls. Dimethyl itaconate injections also increased intracellular itaconate accumulation in PBMCs, by approximately 9-fold, 2.5-fold and 1.5-fold higher than mean values in media, heat-killed MG and live MG-inoculated birds.

**Fig. 1:**
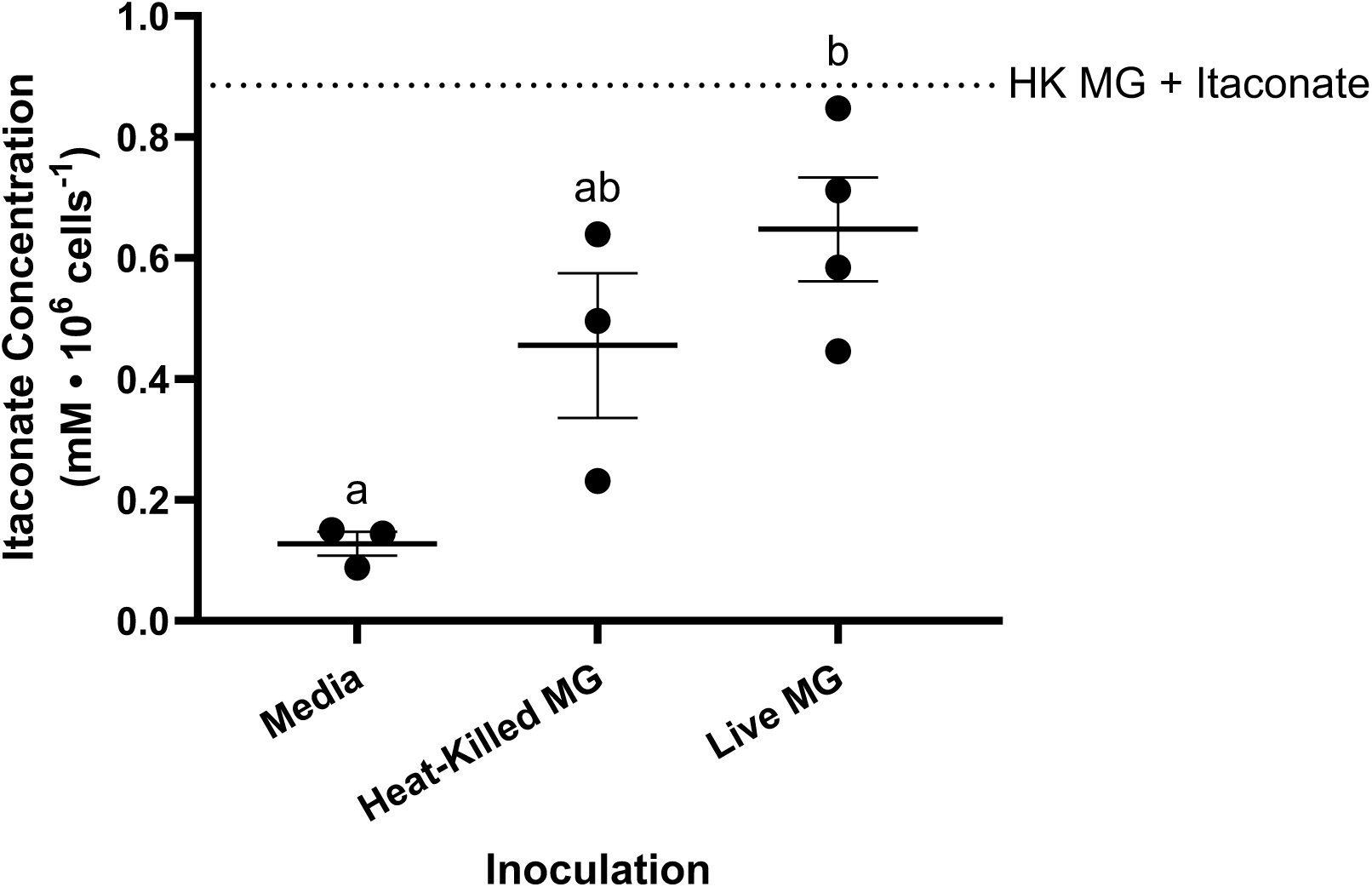
Itaconate accumulation in peripheral blood mononuclear cells from house finches inoculated with *Mycoplasma gallisepticum*. Itaconate concentration standardized to cell count. Dotted line represents itaconate concentration in peripheral blood mononuclear cells from house finches inoculated with heat-killed *Mycoplasma gallisepticum* and intraperitoneally injected with dimethyl itaconate (n = 1). Data plotted as mean ± s.e.m. N = 3-4.

### MG infection suppresses SDH-mediated respiration of immune cells

We found that MG inoculation resulted in significant variation in SDH-mediated OCR in permeabilized PBMCs under State 2 (F_3,20_ = 8.0, p < 0.01), State 3 (F_3,20_ = 4.8, p = 0.01) and State 4o (F_3,20_ = 8.4, p < 0.001) conditions (Fig. 2). Heat-killed MG increased state 2 SDH-mediated OCR, baseline and maximal OCR compared to media controls, suggesting a robust metabolic activation in response to pathogen recognition. In contrast, PBMC OCR was unchanged from media controls during live MG infection, indicating that live MG prevents the metabolic activation normally induced in immune cells following pathogen recognition. In intact PBMCs, the effects of MG on mitochondrial respiration were variable: baseline OCR was marginally affected (W_3,10_ = 3.59, p = 0.05), maximal OCR was significantly affected (F_3,20_ = 7.0, p < 0.01), but leak OCR was unaffected (W_3,9.51_ = 1.58, p = 0.26) (Fig. 3). Specifically, maximal OCR was elevated with heat-killed MG relative to live MG, but both groups were similar to media controls. In contrast, we found little effect of heat-killed or live MG on ECAR (Fig. S3), besides observing low ECAR values during live MG infection relative to other groups. We found no significant differences among groups in baseline ECAR (F_3,19_ = 2.1, p = 0.13) or maximal ECAR (F_3,19_ = 1.7, p = 0.20). These changes in OCR and ECAR elicited by MG also translated to significant variation in total ATP production rates (W_3,7.79_ = 5.68, p = 0.02), nearly significant variation in ATP production from OXPHOS (W_3,8.14_ = 3.20, p = 0.08), but had little effect on ATP production from glycolysis (F_3,18_ = 1.7, p = 0.21) (Fig. S4). We found that heat-killed MG modestly increased (p < 0.1) total ATP production rates relative to live MG but found similar ATP production rates between control birds and live MG-inoculated birds.

**Fig. 2:**
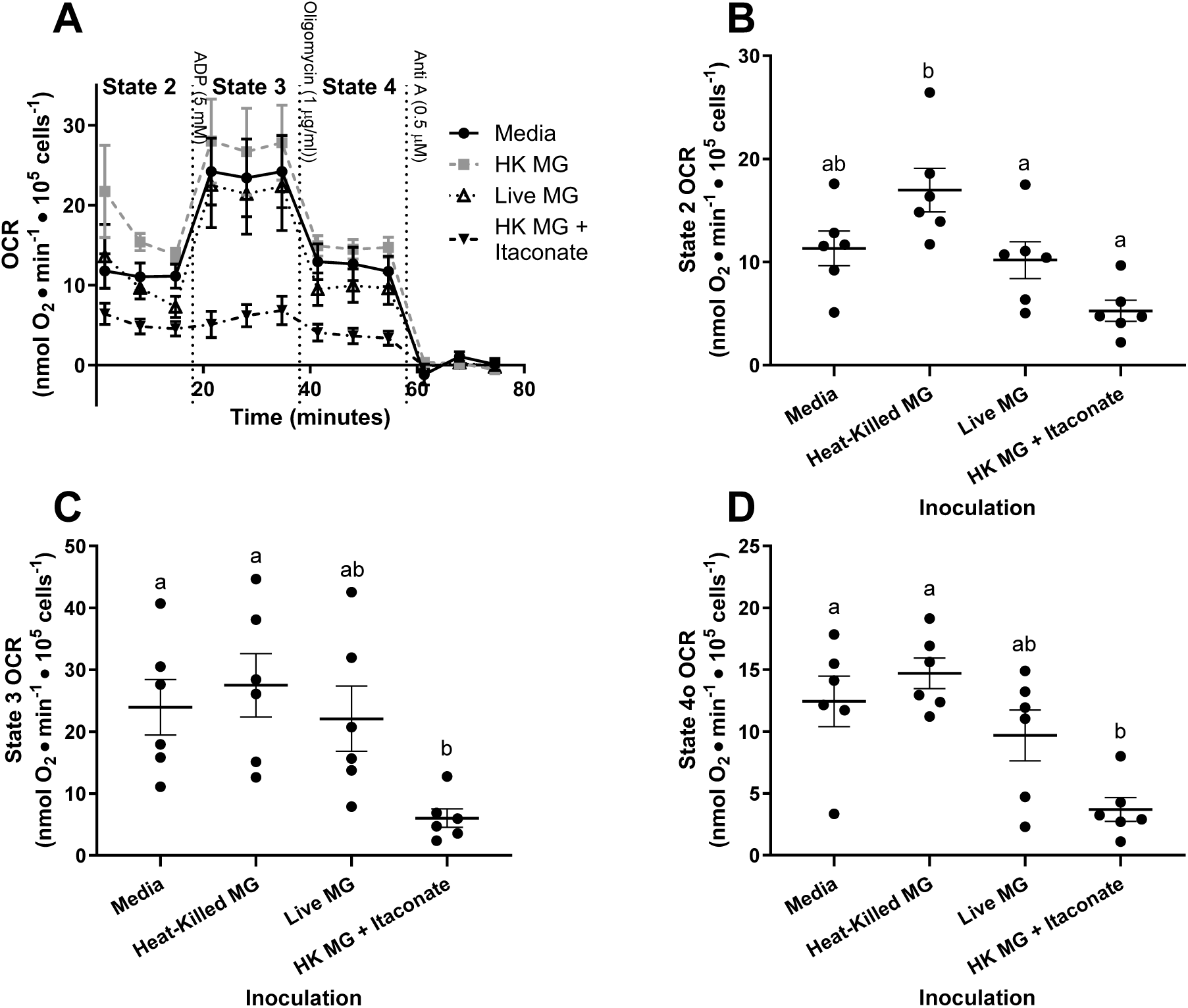
Complex II-mediated O_2_ consumption rate (OCR) in permeabilized peripheral blood mononuclear cells from house finches during acute *Mycoplasma gallisepticum* infection. A) representative trace. B) OCR measured in cells after stimulation of complex II, (C) stimulation of ATP-synthase and D) after inhibition of ATP synthase. OCR standardized to cell count and corrected for non-mitochondrial OCR. Data plotted as mean ± s.e.m. N = 5-6. Significant differences between groups represented by different letters (p < 0.05).

**Fig. 3:**
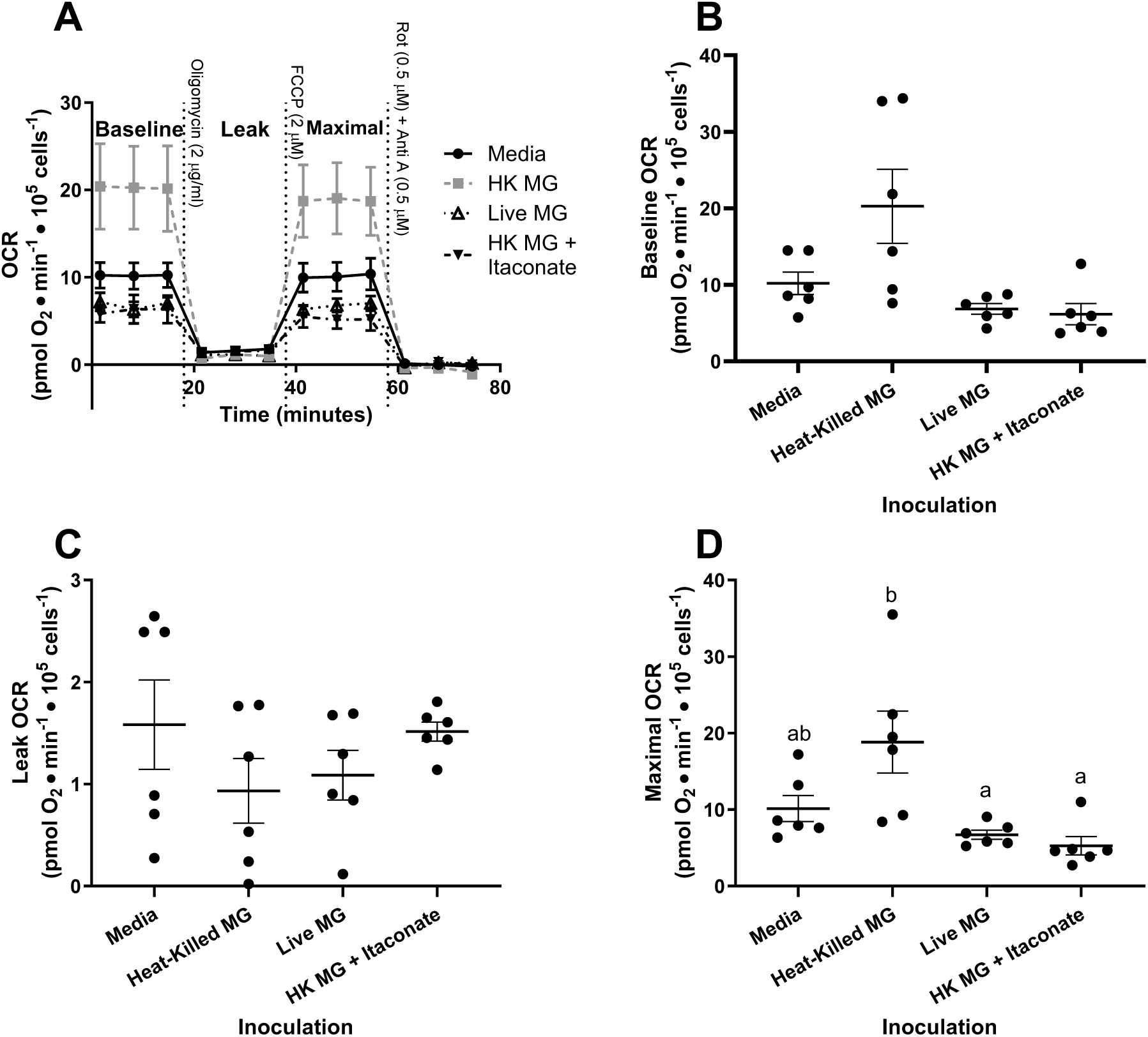
O_2_ consumption rate (OCR) in intact peripheral blood mononuclear cells from house finches during acute *Mycoplasma gallisepticum* infection. A) representative trace. OCR measured in cells without any added substrates or inhibitors (B), after inhibition of ATP synthase (C) and uncoupling of the electron transport system from O_2_ consumption (D). OCR standardized to cell count and corrected for non-mitochondrial OCR. Data plotted as mean ± s.e.m. N = 6. Significant differences between groups represented by different letters (p < 0.05).

### Itaconate mimics live MG

We found that the increased OCR in PBMCs elicited by heat-killed MG was also apparently blocked by itaconate injections (Fig. 2, 3). Itaconate injections elicited decreased SDH-mediated OCR relative to heat-killed MG (p < 0.05), but similar to live MG in all conditions. Itaconate injections also decreased SDH-mediated OCR relative to media control (p < 0.05) in state 3 and state 4o conditions but was similar in state 2. Similarly, itaconate injection was associated with lower maximal OCR than heat-killed MG (p < 0.05) and in both conditions was similar to live MG infection and media control. In contrast, itaconate injection had no detected effect on baseline or leak OCR. We also found that itaconate administration led to lower total ATP production rates than with heat-killed MG (p < 0.1) but was similar to both live MG and media controls (Fig. S3). We did not detect any effect of itaconate administration on glycolytic or OXPHOS ATP production.

### MG blocks inflammatory signaling in erythrocytes

Consistent with apparent activation of PBMC metabolism, heat-killed MG induced strong inflammatory transcriptional responses in erythrocytes (Fig. 6). We found that expression of IL-1β (F_3,19_ = 9.4, p < 0.001), TNFSF15 (W_3,10.2_ = 6.9, p < 0.01) and IL-8L (F_3,19_ = 8.4, p < 0.001) varied with MG inoculation, but the other genes did not: IL-10 (F_3,19_ = 1.5, p = 0.26), IL-18 (F_3,19_ = 0.1, p = 0.94), TGF-β2 (F_3,19_ = 0.3, p = 0.82). In the differentially expressed genes, heat-killed MG upregulated expression relative to media controls, but both live MG and heat-killed MG with itaconate elicited similar expression levels as media controls.

### Tissue mitochondria have differential responses to MG infection

We excluded eyelid OCR measurements in one bird from the heat-killed and itaconate group from our analyses, because our recorded OCR values were at or below instrumental background levels. We investigated the mitochondrial respiration response to MG infection in two infected tissues, the trachea and eyelid conjunctiva, and found little overall effect in the trachea, but a pronounced response in the eyelid (Fig. 4; Fig. 5). MG infection did not affect SDH-mediated OCR in trachea in state 3 (F_3,20_ = 0.91; p = 0.45) or state 4o conditions (F_3,20_ = 3.06; p = 0.05), but live MG infection increased SDH-mediated OCR in eyelid under state 3 by approximately 3-fold relative to media controls, (F_3,19_ = 4.42; p = 0.02) but had little effect on state 4o measurements (F_3,19_ = 2.53; p = 0.09). Similarly, live MG had little effect on complex I-mediated respiration in state 2 (F_3,20_ = 4.41; p = 0.02) and state 3 (F_3,20_ = 2.03; p = 0.14) conditions in the trachea, but increased state 2 (F_3,19_ = 27.31; p < 0.0001) and state 3 (F_3,19_ = 10.20; p < 0.001) OCR in the eyelid by nearly 3-fold. We also found tissue-specific variation in non-mitochondrial OCR (Fig. S5), where MG again had little effect in the trachea (F_3,20_ = 4.30; p = 0.02), but heat-killed MG and live MG increased non-mitochondrial OCR in the eyelid (F_3,19_ = 18.31; p < 0.0001) by c. 2-fold and 4-fold, respectively. Itaconate injections had little effect on mitochondrial and non-mitochondrial OCR in either tissue and were consistently similar to media controls and heat-killed MG.

**Fig. 4:**
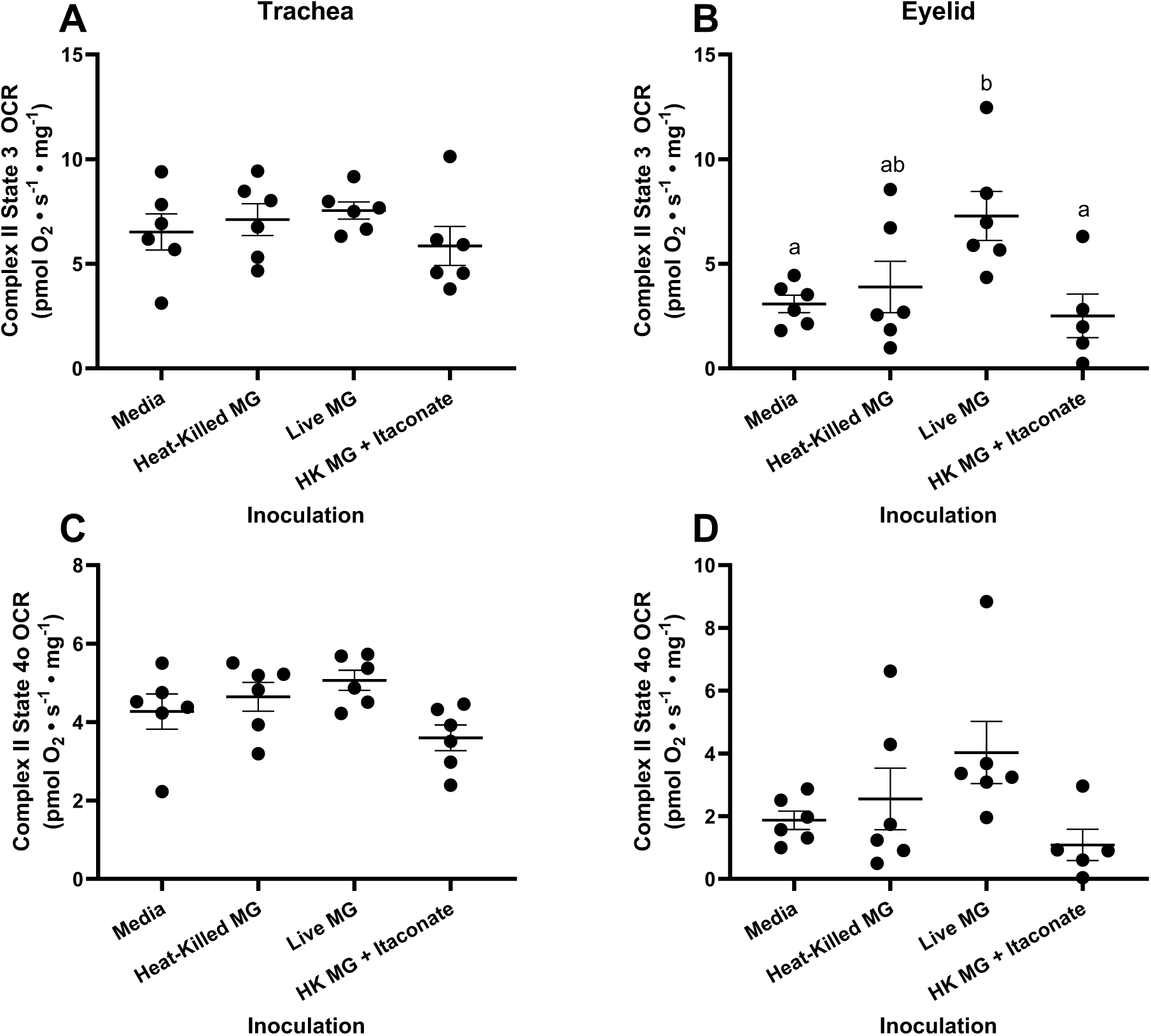
Succinate dehydrogenase-mediated O_2_ consumption rate (OCR) in permeabilized tissues from house finches during acute *Mycoplasma gallisepticum* infection. OCR measured in trachea (A, C) and eyelid conjunctiva (B, D). OCR measured with maximal stimulation of complex II and ATP synthase (A, B) and following inhibition of ATP synthase (C, D). OCR standardized to tissue wet mass and corrected for non-mitochondrial OCR. Data plotted as mean ± s.e.m. N = 5-6. Significant differences between groups represented by different letters (p < 0.05).

**Fig. 5:**
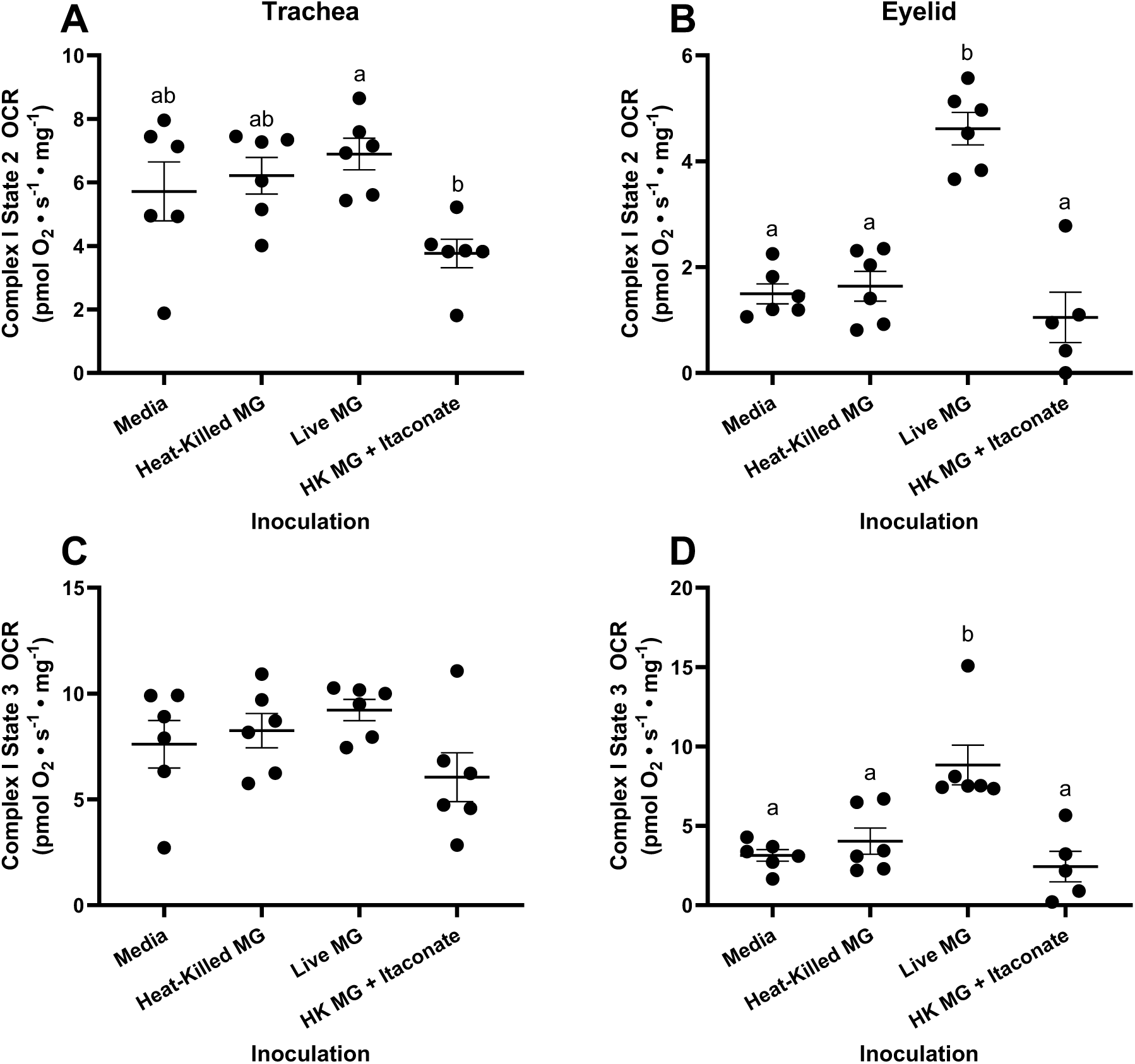
Complex I-mediated O_2_ consumption rate (OCR) in permeabilized tissues from house finches during acute *Mycoplasma gallisepticum* infection. OCR measured in trachea (A, C) and eyelid conjunctiva (B, D). OCR measured with maximal stimulation of complex I (A, B) and following maximal stimulation of ATP synthase (C, D). Values standardized to tissue wet mass and corrected for non-mitochondrial OCR. Data plotted as mean ± s.e.m. N = 5-6. Significant differences between groups represented by different letters (p < 0.05).

**Figure 6:**
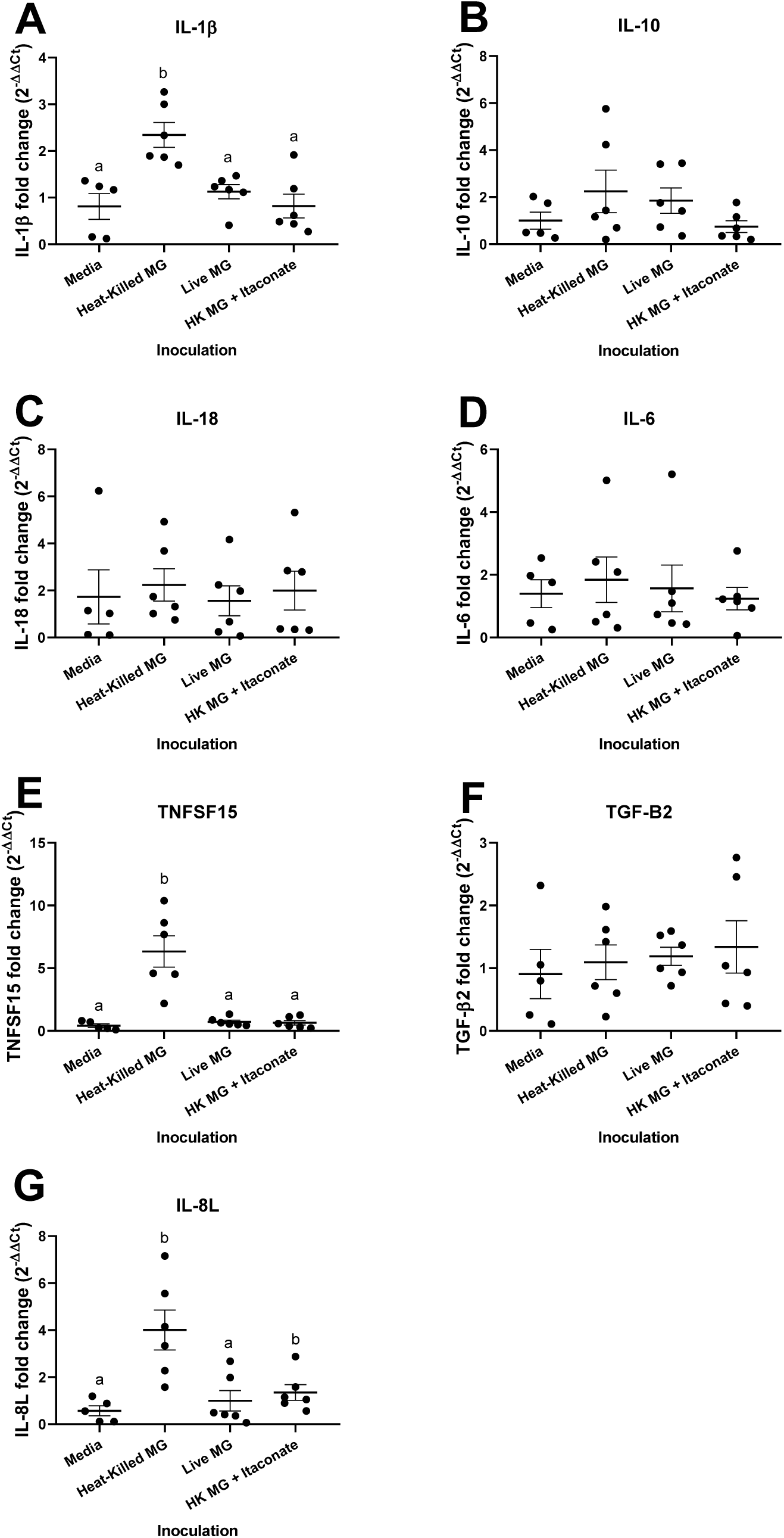
Expression of immune response-related genes in erythrocytes from house finches during acute *Mycoplasma gallisepticum* infection. Expression of interleukin-1β (A), interleukin-6 (B), interleukin-8L (C), interleukin-10 (D), interleukin-18 (E), TGF-β2 (F) and TNFSF15 (G). Gene expression expressed 2^-ΔΔCt^ relative to 28S rRNA in the media control group. Data plotted as mean ± s.e.m. N = 5-6. Significant differences between groups represented by different letters (p < 0.05).

### MG upregulates pro-inflammatory gene expression in eyelid

Despite suppressed immune cell metabolism and cytokine transcription in peripheral erythrocytes, live MG induced strong local inflammatory transcriptional responses. We found that expression of all immune genes significantly varied with inoculation (Fig. 7): IL-10 (W_3,10.4_ = 11.9, p < 0.01) IL-18 (F_3,20_ = 3.5, p = 0.03), IL-1β (W_3,10.0_ = 11.3, p < 0.01), IL-6 (F_3,20_ = 7.9, p < 0.01), IL-8L (W_3,10.0_ = 11.1, p < 0.01), TGF-β2 (F_3,20_ = 4.1, p = 0.02) and TNF-SF15 (F_3,20_ = 16.6, p < 0.001). Live MG induced upregulation of all genes except for TGF-β2 and IL-18 relative to media controls, while heat-killed MG and itaconate injections had no effect on expression for any gene.

**Figure 7:**
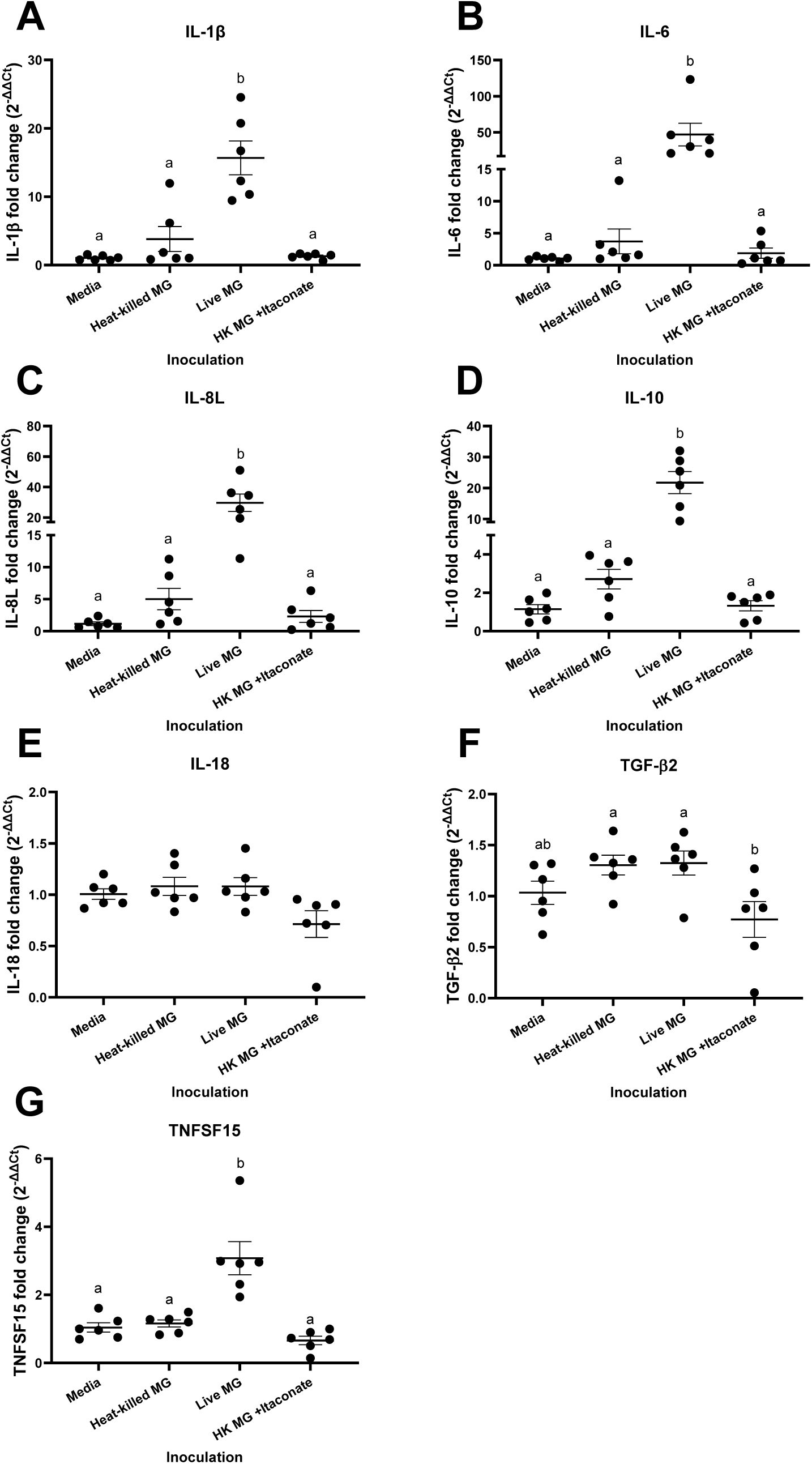
Expression of immune response-related genes in eyelid conjunctiva from house finches during acute *Mycoplasma gallisepticum* infection. Expression of interleukin-1β (A), interleukin-6 (B), interleukin-8L (C), interleukin-10 (D), interleukin-18 (E), TGF-β2 (F) and TNFSF15 (G). Gene expression expressed 2^-ΔΔCt^ relative to 28S rRNA in the media control group. Data plotted as mean ± s.e.m. N = 5-6. Significant differences between groups represented by different letters (p < 0.05).

## Discussion

In this study, we assessed the mechanisms underlying the suppressive effects of MG infection on host immunometabolism in a wild songbird. We tested the hypotheses that 1) MG host mitochondrial metabolism suppression is systemic and 2) this suppression acts through a mechanism mediated by the immunomodulatory metabolite itaconate. Contrary to our hypothesis 1, live MG increased mitochondrial respiration and pro-inflammatory cytokine gene expression in the eyelid conjunctiva but had little effect in trachea. In contrast, live MG suppressed mitochondrial respiration in PBMCs and pro-inflammatory cytokine gene expression in RBCs. In accordance with our hypothesis 2, we found that exogenous dimethyl itaconate administration produced a similar attenuating effect on mitochondrial respiration in PBMCs and cytokine gene expression in RBCs compared with live MG infection. Together, our results demonstrate that the host immunometabolic responses to MG are highly tissue specific and that MG specifically suppresses immunometabolic responses in circulating immune cells during the early stages of infection in a similar mechanism as itaconate.

### Pathogen suppression of early systemic immune metabolic responses

Host immunometabolic changes are necessary to both induce the signaling that initiates the innate immune response and to meet increased ATP demands of immune cells during infection [4,54]. In the present study, we observed that live MG blunted the increase in PBMC mitochondrial respiration induced by heat-killed MG. More importantly, the reduction in mitochondrial respiration observed in PBMCs from live MG-infected birds was largely attributable to suppressed SDH-mediated respiration (Figs. 2B and 3B). Alterations in mitochondrial respiratory chain function play a critical role in regulating innate immune responses, particularly during antibacterial defense in macrophages [55], monocytes [56], neutrophils [57] and dendritic cells [58]. Within the mitochondrial respiratory chain, SDH serves as a key metabolic hub that links cellular metabolism to immune activation and inflammatory signaling during innate immune responses [59]. Therefore, the apparent metabolic suppression observed in immune cells during live MG infection in this study suggests that MG disrupts the metabolic changes necessary to mount an innate immune response.

Avian erythrocytes are dynamic, nucleated cells that retain their full complement of organelles [60]. Accumulating evidence suggests that these cells participate in immune-related processes [52,61], so we used erythrocyte gene expression as a proxy for systemic immune responses. Moreover, previous studies have shown that MG can invade chicken RBCs both *in vitro* and *in vivo* [62]. Indeed, heat-killed MG significantly increased expression of IL-1β, TNFSF15, and CXCLi2 (the avian homolog of IL-8) in erythrocytes (Fig. 6), but these changes were prevented by live MG. This finding is consistent with previous work showing minimal changes in blood cytokine levels during the early stages of MG infection in house finches [51]. These pro-inflammatory cytokines are regulated, at least in part, through SDH-dependent pathways involving hypoxia-inducible factor 1α (HIF-1α), linking mitochondrial metabolism to immune activation [59,63,64]. This pattern is consistent with our metabolic findings and further indicates that MG exerts a suppressive effect on immunometabolism in circulating immune cells during the early stages of infection.

Upon pathogen recognition, immune cells typically undergo metabolic reprogramming characterized by a shift from oxidative phosphorylation toward aerobic glycolysis, a phenomenon similar to the Warburg effect [6,65]. Indeed, enhanced glycolytic activity has been shown to promote host defense and pathogen clearance during infections caused by several Mycoplasma species [66,67]. Several pathogens have evolved mechanisms to inhibit glycolytic reprogramming in host immune cells, thereby limiting effective inflammatory responses [10,11,68,69]. However, we did not detect significant differences in glycolytic rates among treatment groups but noted a potential decrease in glycolytic rates in birds inoculated with live MG (Fig. S3). The metabolic transition in immune cells typically elicited during infection supports the rapid ATP production and biosynthetic demands required for inflammatory responses [5]. Here, we found that total ATP production rates in PBMCs were elevated with heat-killed MG (Fig. S4), likely reflecting the elevated energetic demands associated with immune activation, including cytokine synthesis and secretion [70]. Interestingly, the increased ATP production rate was blocked by live MG inoculation. These patterns in ATP production rate in PBMCs were mirrored by SDH-mediated respiration and expression of pro-inflammatory cytokines in RBCs. Our findings indicate that the suppressive effects of MG and itaconate on immune cell metabolism may translate into reductions in supplying the ATP necessary for immune activation.

### Itaconate as a mediator of pathogen tolerance

We found that live MG inoculation increased itaconate levels in PBMCs of house finches. Moreover, heat-killed MG combined with dimethyl itaconate treatment produced inhibitory effects similar to live MG on mitochondrial respiration in PBMCs and pro-inflammatory cytokine gene expression in RBCs, suggesting that the immunometabolic suppressive effects of MG may operate through mechanisms involving itaconate [40]. Consistent with this, previous studies have shown that elevated itaconate can be detrimental to the host during infection with other *Mycoplasma* species [39,40], as well as during infections by diverse bacterial and viral pathogens [71–75].

MG-induced increases in host itaconate levels may promote progression toward a disease tolerance state [43,47]. MG causes chronic respiratory disease in avian species, and studies examining temporal invasion dynamics of MG across house finch populations have shown reduced upregulation of pro-inflammatory genes in populations with a longer history of MG exposure compared with populations experiencing more recent pathogen endemism [43]. For example, Alabama birds, which have a substantially longer history of MG exposure, exhibited greater tolerance, characterized by reduced pathology and pathogen load, lower IL-1β expression, and higher IL-10 levels compared with Arizona birds during early stages of infection [47]. All of these are tightly regulated by SDH in host immune cells. Behavioral tolerance is similarly increased in house finch populations with longer MG exposure [76], but whether this aspect of pathogen tolerance is linked to variation in SDH signaling remains unclear. Together, these findings suggest that the evolution of increased infection tolerance in house finches following the emergence of MG may partly result from the immunometabolic suppressive properties of MG, potentially through itaconate-mediated inhibition of host mitochondrial SDH.

The mechanisms by which MG exploits itaconate to suppress host immunometabolism remain largely unresolved and is beyond the scope of this study. The MG genome does not encode any obvious proteins capable of synthesizing itaconate [77], so manipulation of host anabolic pathways appears likely. Indeed, previous studies have demonstrated that Mycoplasma infection alters multiple tricarboxylic acid (TCA) cycle metabolites in host cells, including succinate, itaconate, isocitrate, malate, and oxaloacetate [39,78]. One potential mechanism for MG to stimulate itaconate synthesis in host cells is to induce upregulation of the gene encoding the itaconate-synthesizing enzyme (*Irg1*) following NF-κB- and STAT1-mediated signaling downstream of TLR2 activation [40]. An underexplored alternative (but non mutually exclusive) mechanism is that MG directly synthesizes itaconate. Mycoplasma species exhibit limited capacity to synthesize TCA cycle metabolites (e.g. fumarate, malate) [77], potentially via anaplerotic reactions independent of conventional TCA cycle enzymes [79]. This suggests that alternative enzymes may be responsible for synthesizing some TCA cycle metabolites [80,81]. In summary, it is likely that MG-induced alterations of host cell metabolism drive the increase in intracellular itaconate levels observed in PBMCs of live MG-infected birds. However, it is also possible that MG itself produces TCA cycle metabolites, or their precursors, that contribute to initiating this process. Both processes should receive attention in future investigations.

### Tissue-specific metabolic patterns during infection

We observed marked differences in metabolic responses to MG infection between eyelid conjunctiva and PBMCs. At the primary site of infection, high pathogen loads lead to infiltration of lymphocytes, heterophils, and plasma cells in the eyelid conjunctiva [82,83]. We conducted our measurements at 3 days post inoculation, a time point when conjunctivitis was first observed in previous studies [84], suggesting that immune cell infiltration had already occurred at this stage. Because respiration measurements in permeabilized conjunctiva could not distinguish mitochondrial activity among different cell types, we speculate that the observed increase in both mitochondrial and non-mitochondrial respiration following live MG inoculation reflects the high ATP demands associated with pro-inflammatory cytokine production, as well as elevated NADPH oxidase activity required for respiratory burst in infiltrating immune cells [85,86]. We predict that a similar metabolic response may also occur in the trachea under conditions where tracheal lesions and immune cell infiltration are present, such as during later stages of infection [87–89] or following airway inoculation [90]. Consistent with this interpretation, we found that the majority of inflammatory cytokine genes measured were upregulated in eyelid conjunctiva following live MG inoculation. This agrees with previous studies showing that eyelid conjunctiva exhibits elevated inflammatory cytokine expression between days 3 and 6 post inoculation [51].

Further, this dichotomy between immune/metabolic suppression in local vs. peripheral tissues likely holds advantages for the pathogen. Recent work in this system has shown that more severe local pathology (i.e., conjunctivitis) results in a higher amount of pathogen shedding onto the surface of bird feeders and more efficient transmission to conspecifics [91–94]. As such, the increased inflammatory signaling and metabolic responses at the local site of infection we observed are likely beneficial to the pathogen in terms of onward transmission. On the other hand, suppression of systemic immune responsiveness, as evidenced here by lower cytokine transcription and mitochondrial respiration in peripheral blood cells under live-MG treatment, should benefit the pathogen by muting activation of subsequent immune responses, including adaptive immune defenses that appear most effective against this pathogen [14,95,96].

### Limitations

We acknowledge that our study has several limitations. While the use of wild-caught house finches provides the advantages of greater ecological relevance and a more realistic representation of host-pathogen interactions, the species remains a non-model organism, which constrains the range of available methodologies. For example, because our study species lacks commercially available antibodies for immune markers, has limited plasma proteomic resources, and yields relatively low numbers of PBMCs from small blood samples, we were unable to directly assess the detailed mechanisms underlying itaconate synthesis and its mode of action [40]. We suggest that the experimental design used here could be repeated with immune cells *in vitro*, such as with bone marrow-derived macrophages isolated from chickens, which could be a promising avenue to further study immunometabolic suppression by MG. Similarly, we acknowledge that while administration of exogenous dimethyl itaconate is an effective way to increase intracellular itaconate levels in *in vivo* systems, this itaconate analogue can induce artefactual effects on cellular physiology that are distinct from endogenously produced itaconate [44]. For example, dimethyl itaconate has greater electrophilic strength than itaconic acid, which can lead to post-translational modification of immune-related proteins, or increased reactive oxygen species production following glutathione depletion [97,98]. Future studies could address this limitation with itaconic acid supplementation of bone marrow-derived macrophages *in vitro* [98].

### Implications

As a recently emerged and well-documented host–pathogen system, the house finch–MG system provides an exceptional opportunity to identify evolutionary patterns underlying host–pathogen interactions [99,100]. Past studies have shown that, across house finch populations, longer exposure history to MG is associated with reduced inflammatory signaling, indicating a greater propensity for tolerance to MG infection [43,47,76,85]. At the same time, more evolutionarily derived MG strains have become increasingly virulent by inducing stronger host inflammatory responses and greater pathology [85,101,102]. Our data suggest that these evolutionary adaptations may be partly mediated by the immunometabolic properties of MG, potentially through targeting host mitochondrial SDH.

Similarly, little is known about how MG underwent a major host shift from domesticated poultry to wild house finches and evolved the ability to induce severe pathology in this species, but not in other closely related wild bird species [103–105] that share similar transmission routes [106]. We hypothesize that the ability to manipulate host immunometabolism may represent a key factor contributing to MG host specificity, its successful emergence in the novel house finch host, and subsequent and long-term endemism. Future studies examining whether MG strains with differing virulence vary in their effects on itaconate/SDH-regulated pathways, especially if such effects differ at local vs systemic levels, as well as comparative studies between MG-susceptible and MG-resistant avian species, could provide important insight into the role of immunometabolism in shaping host susceptibility and the evolutionary dynamics of host–pathogen interactions.

MG is among the most economically devastating respiratory pathogens affecting the global poultry industry [107]. Current control strategies are largely limited to biosurveillance measures, along with live attenuated MG vaccines [109]. Consequently, the persistent nature of MG infection highlights the need for alternative therapeutic approaches. Our findings suggest that the itaconate/SDH axis is a potential therapeutic target for MG control. Pharmacological agents such as citraconate, which inhibits the ACOD1/*Irg1*-mediated itaconate synthesis pathway, may offer potential therapeutic value for treating MG infections in birds [40,110]. Future studies are needed to evaluate whether targeting the itaconate/SDH pathway can provide an effective therapeutic strategy for controlling MG infection in poultry.

## Conclusions

Here, we show that MG suppresses metabolic reprogramming of immune cells in house finches, likely by targeting mitochondrial SDH through a mechanism resembling itaconate-mediated metabolic regulation. Our findings add to growing evidence that certain pathogens have evolved mechanisms to manipulate host metabolic pathways that underlie innate immune responses, ultimately to the detriment of the host. More broadly, our results identify immunometabolic suppression as a bacterial strategy for modulating host defense, highlighting mitochondria as a critical battleground in the evolutionary arms race between hosts and pathogens.

## Acknowledgements

We would like to thank Dr. Dana Hawley (Virginia Tech) for capturing the birds used in this study. We would like to thank the veterinary staff at the University of Memphis for providing care to the birds used in this study. We would also like to thank Stefane Saruhashi (Cornell U) for providing editorial comments on this manuscript.

## Data availability

All data and Western Blot images are publicly available at FigShare (DOI 10.6084/m9.figshare.33167168).

