## Supplementary materials for "*Mycoplasma gallisepticum* uses itaconate-associated mitochondrial inhibition to suppress host immunometabolism"

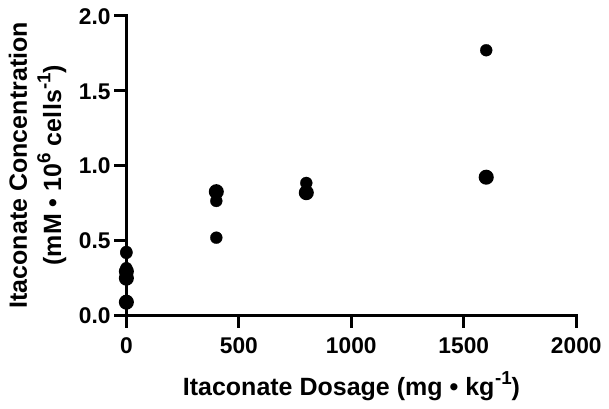

**Figure S1**: **Dose response of exogenous dimethyl itaconate administration and intracellular itaconate concentrations in peripheral blood mononuclear cells in house finches.** Dimethyl itaconate administered via intraperitoneal injections and dosage is expressed relative to body mass.

*Western blots*

To assess the effectiveness of our tissue and PBMC permeabilization prior to respirometry experiments, we used Western blotting to measure the abundance of cytochrome c in PBMCs, trachea and eyelids with and without detergent digestion. From 2 birds, we dissected the trachea and eyelid conjunctivae and divided the tissue for each bird in two. We digested one half of each tissue for each bird with saponin as described in the Mitochondrial Experiments section and the other half was incubated in ice-cold isolation solution for 20 minutes, then samples were frozen at -80°C until further analyses. We also prepared 2 PBMC extracts pooled from 6 birds each. We digested one PBMC extract with digitonin while the other was incubated for similar time in Seahorse media. We homogenized the samples in lysis buffer (RIPA buffer with 0.8 mM DTT, 0.4 mM PMSF, 1x phosphatase and 1x protease inhibitor), using a BeadBlaster at 4000 rpm for 2 cycles of 30 seconds 4°C, followed by 3 cycles of sonication for 10 seconds on ice and centrifugation at 16,000 g for 15 minutes at 4°C. We removed the supernatant and measured protein content with a BCA assay. We then mixed the samples with 2x sample buffer (4% SDS, 20% glycerol, 5% β-mercaptoethanol, 2 mM EDTA, 0.1 mg mL^-1^ bromophenol blue, 100 mM Tris-HCl, pH 6.8) and denatured proteins via incubation at 100°C for 5 minutes followed by cooling at room temperature for 5 minutes.

We then conducted SDS-PAGE, loading 5-10 µg of protein per well for tissue and comparing with 2 µL of protein ladder (Bio-Rad 1610373). On a separate gel, we loaded 5 µg of protein for PBMCs and protein ladder. We used 10% polyacrylamide gels and ran the gel at 110V for 45-60 minutes in running buffer (in g L^-1^: 3.03 Trizma base, 14.4 glycine, 1 SDS). Following band separation, we transferred to a nitrocellulose membrane using a Trans-Blot Turbo (Bio-Rad), using a proprietary transfer buffer (Bio-Rad 10026938). We cut the membrane at the 25 kDa ladder band to GAPDH (c. 36 kDa) and cytochrome c (c. 12 kDa). We blocked the membranes with 1 hour incubation at room temperature in skim milk dissolved in TBS-T (0.05 g mL^-1^). We washed the membranes 3 times with TBS-T, immersed them in rabbit primary antibody diluted in blocking buffer and incubated at 4°C overnight. Cytochrome c was diluted to 1:200 (GeneTex GTX02626) and GAPDH was diluted 1:1000 (GeneTex GTX100118). We then rinsed the membranes with milliQ water 4 times, followed by 3 washes with TBS-T, each for 10 minutes. We then added HRP-conjugated goat anti-rabbit secondary antibody (Abclonal AS014) diluted 1:1000 in TBS-T. We repeated TBS-T washing and incubated membranes with ECL substrate (Thermo 34580) before imaging chemiluminescence using an iBright 1500 (Invitrogen).

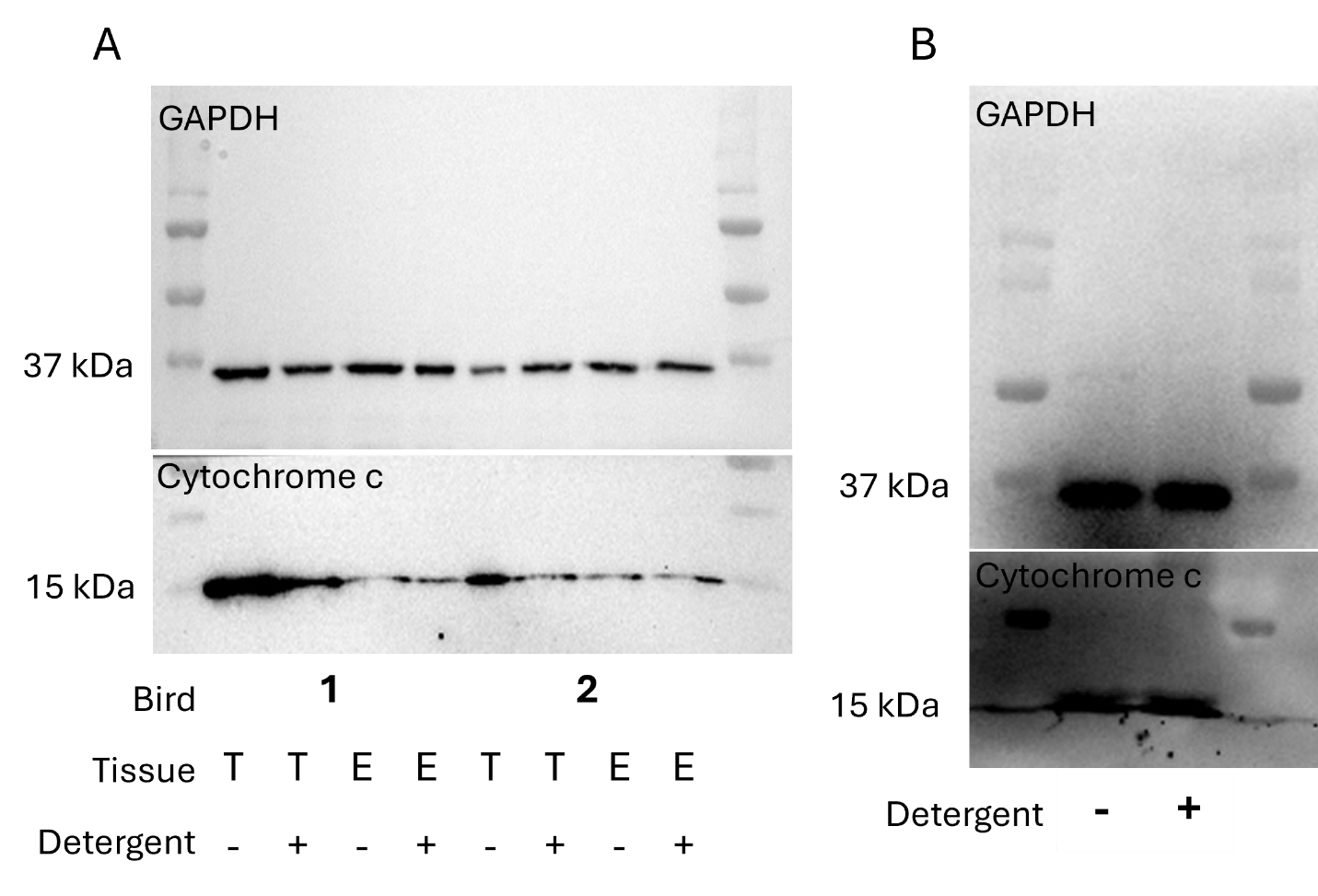

**Figure S2**: **Western blots for cytochrome c and glycerol-3-phosphate dehydrogenase (GAPDH) in tissues (A) and peripheral blood mononuclear cells (B).** A) trachea (T) and eyelid conjunctivae (E) samples were incubated with or without saponin, repeated for two individual birds. B) cells were incubated with or without digitonin.

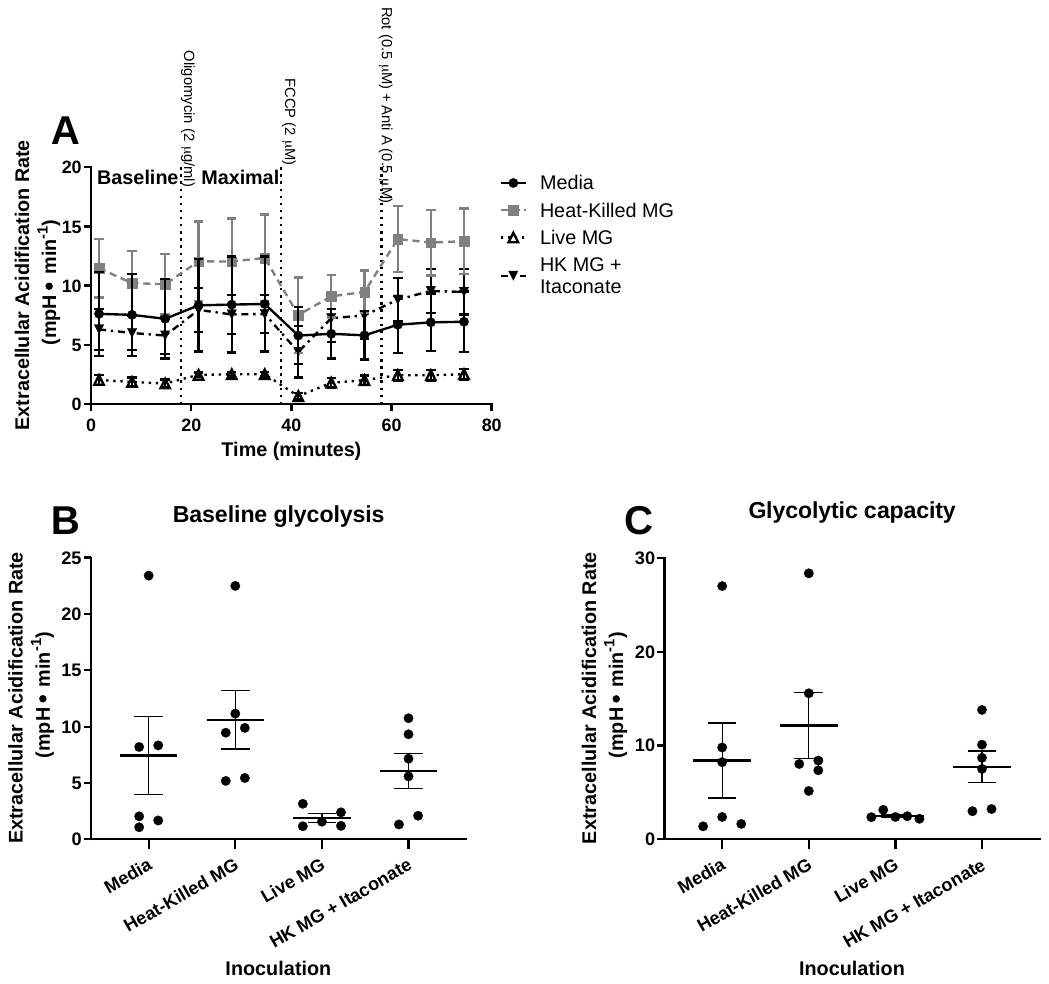

**Figure S3: Extracellular acidification rate (ECAR) in intact peripheral blood mononuclear cells from house finches during acute *Mycoplasma gallisepticum*** **infection**. (A) representative trace. ECAR measured in cells without any added substrates or inhibitors, representing baseline glycolysis. (B) after inhibition of ATP synthase, representing glycolytic capacity. Data plotted as mean ± s.e.m. N = 6.

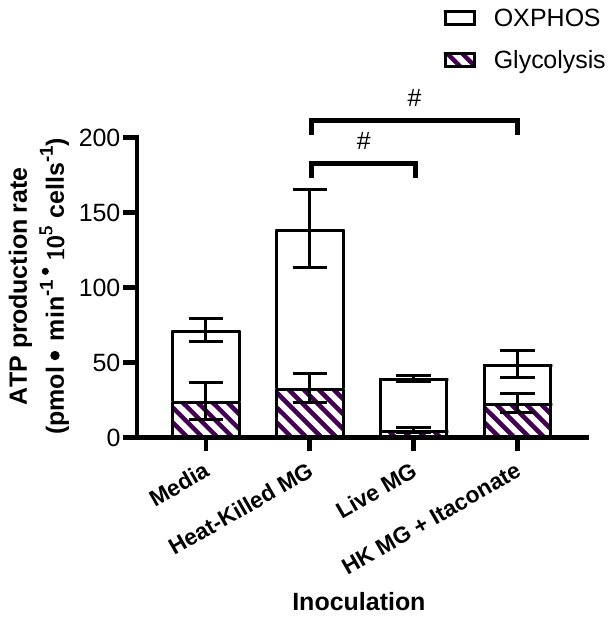

**Figure S4: ATP production rates in intact peripheral blood mononuclear cells from house finches during acute *Mycoplasma gallisepticum*** **infection**. ATP production rates calculated using oxygen consumption rate (OXPHOS – empty columns) and extracellular acidification rate (glycolysis – hatched columns). ATP production rates standardized to cell count. # represent differences approaching significance (p < 0.1) among groups in total ATP production rate. Data plotted as mean ± s.e.m. N = 5-6.

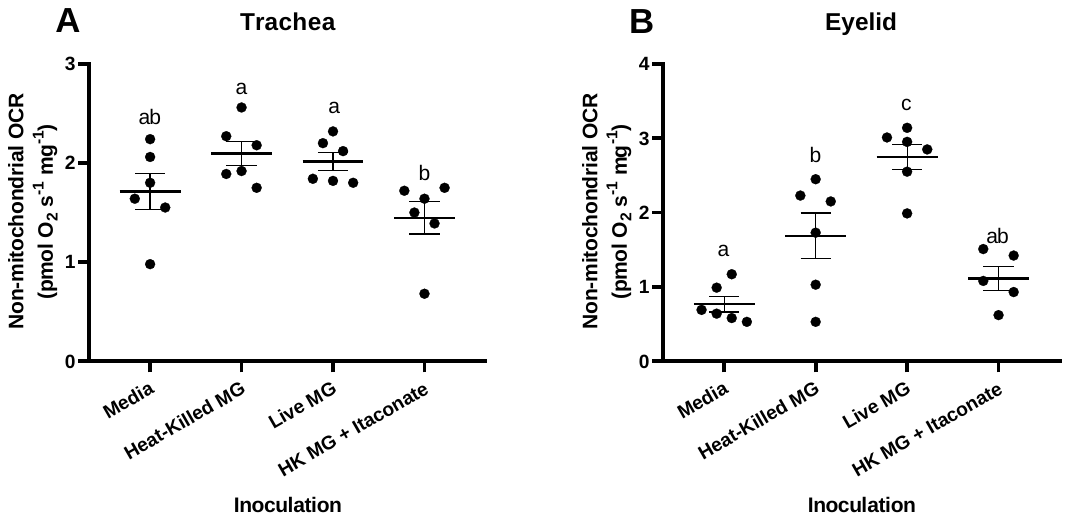

**Figure S5: Non-mitochondrial O_2_ consumption rate (OCR) in permeabilized tissues from house finches during acute *Mycoplasma gallisepticum*** **infection**. OCR measured following inhibition of the mitochondrial electron transport system in trachea (A) and eyelid conjunctiva (B). Values standardized to tissue wet mass. Data plotted as mean ± s.e.m. N = 5-6. Significant differences between groups represented by different letters (p < 0.05).

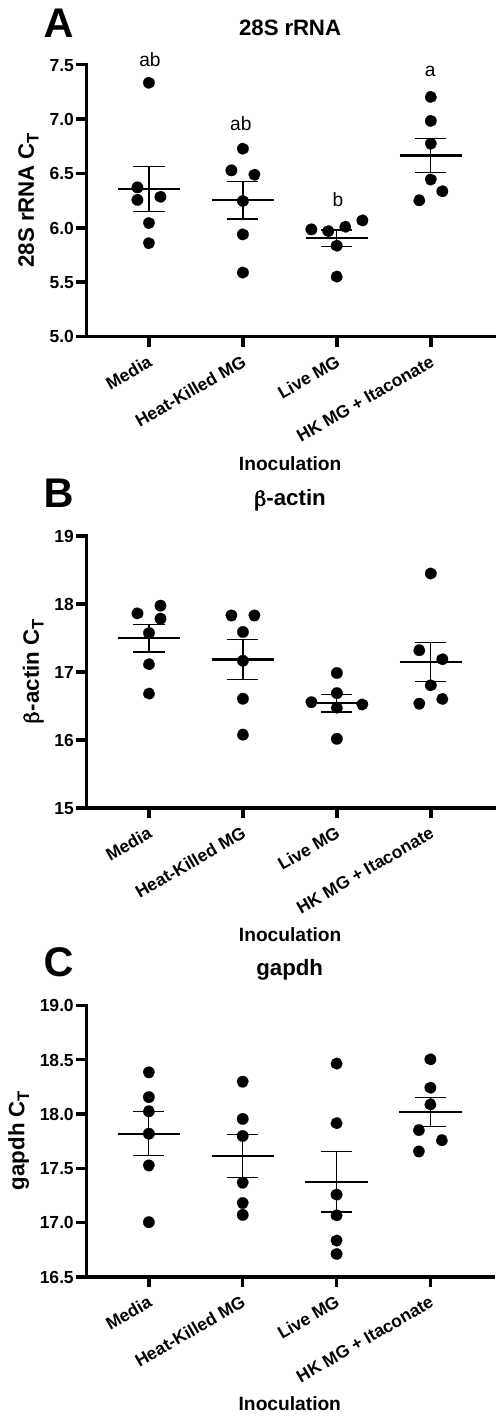

**Figure S6:** **Expression of reference genes in eyelid conjunctiva from house finches during acute *Mycoplasma gallisepticum*** **infection.** Expression of *28SrRNA* (A), *ACTB* (B) and *GAPDH* (C). Gene expression expressed as C_T_. Data plotted as mean ± s.e.m. N = 5-6. Significant differences between groups represented by different letters (p < 0.05).

**Table S1: Primers used for qPCR experiments.**

| Target gene | Primer sequence (5’ to 3’) | |
| --- | --- | --- |
| *IL1B* | Forward  Reverse | TGCTGGACAGAAAGTGAAGCT  GCTGGTAGCCCTTGATGC |
| *IL6* | Forward  Reverse | CAGCGAAAACCAAAATGTTG  GTGTGGAGTGATTCCTGG |
| *IL8L* | Forward  Reverse | CAGTGCATAGCCACTCATTC  GCACACCTCTTTGCCATTC |
| *IL10* | Forward  Reverse | AACCTCCTGCTGAACCTG  ATGTGCTCCATGCTCCTG |
| *IL18* | Forward  Reverse | TCAGTGTCCAGGTGGAAAAC  CTCCTTCCTTGAACCTCACG |
| *TGFB2* | Forward  Reverse | GGCTCCATCACAGAGACAGG  TCTTGCTTCAAGCTCCTCAC |
| *TNFSF15* | Forward  Reverse | GGGGCTCTCACTTTCTG  AGGTCCTGCCTCTTCAC |
| *28SrRNA* | Forward  Reverse | GGCGAAGCCAGAGGAAACT  GACGACCGATTTGCACGTC |
| *GAPDH* | Forward  Reverse | CATCCTGGCATACACAGAG  GTCGTTCAGTGCAATGCC |
| *ACTB* | Forward  Reverse | CATTGCTGACAGGATGCAG  CCGATCCAGACAGAGTATTTG |
